# Genome-wide regulatory effects of STRs stabilized by elevated expression of antioxidant genes in *C. elegans*

**DOI:** 10.1101/2022.09.27.509703

**Authors:** Gaotian Zhang, Erik C. Andersen

## Abstract

Genetic variation can cause significant differences in gene expression among individuals. Although quantitative genetic mapping techniques provide ways to identify genome-wide regulatory loci, they almost entirely focus on single nucleotide variants (SNVs). Short tandem repeats (STRs) represent a large source of genetic variation with potential regulatory effects. Here, we leverage the recently generated expression and STR variation data among wild *Caenorhabditis elegans* strains to conduct a genome-wide analysis of how STRs affect gene expression variation. We identify thousands of expression STRs (eSTRs) showing regulatory effects and demonstrate that they explain missing heritability beyond SNV-based expression quantitative trait loci. We illustrate specific regulatory mechanisms such as how eSTRs affect splicing sites and alternative splicing efficiency. We also show that differential expression of antioxidant genes might affect STR variation systematically. Overall, we reveal the interplay between STRs and gene expression variation in a tractable model system to ultimately associate STR variation with differences in complex traits.

## Introduction

Genetic variation can cause significant differences in gene expression among individuals. Mutations in regulatory elements, such as promoters and enhancers, might only affect the expression of single genes, whereas mutations altering structures and abundances of diffusible factors, such as transcription factors (TFs) and chromatin cofactors, might affect the expression of multiple genes across the genome. Quantitative genetic mapping techniques, including both linkage and genome-wide association (GWA) mapping studies, enable the identification of genome-wide variants that influence gene expression and other complex traits. A genomic locus that contains alleles showing significant association with mRNA expression variation is called an expression quantitative trait locus (eQTL)^1–5^. Although thousands of eQTL have been detected in different organisms, associated genetic variants are mostly limited to single nucleotide variants (SNVs) and short insertions or deletions (indels)^1–11^. Emerging studies successfully linked gene expression variation to other types of DNA variants, such as short tandem repeats (STRs) and structural variants^12–19^.

STRs are repetitive elements consisting of 1-6 bp DNA sequence motifs^17,20^. Compared to SNVs and short indels, STR mutations show 1) orders of magnitude higher mutation rates^20–23^, 2) higher incidence of insertions or deletions, mostly in the number of repeats^24,25^, 3) more multiallelic sites^26^, and 4) more *de novo* mutations^20,26^. Dozens of human diseases have been associated with STR mutations^24^. Various effects of STR variation on regulation of gene expression have also been suggested from both *in vitro* and *in vivo* studies across a wide range of taxa^27–33^. However, these STRs only represented a small fraction of STRs in genomes. To our best knowledge, systematic evaluation of genome-wide associations between STR variation and gene expression variation have only been applied in humans^12,17,34^ and *Arabidopsis thaliana^16,19^*, in part because of the difficulties in accurately genotyping STRs throughout the genome in large scales^35^.

We have recently studied the natural variation in gene expression^5^ and STRs^36^ across wild strains of the nematode *Caenorhabditis elegans*. We collected reliable expression measurements for 25,849 transcripts of 16,094 genes in 207 *C. elegans* strains using bulk mRNA sequencing and identified 6,545 eQTL underlying expression variation of 5,291 transcripts of 4,520 genes using GWA mappings^5^. We characterized 9,691 polymorphic STRs (pSTRs) with motif lengths of 1-6 bp across the species, including the 207 strains above, using high-throughput genome sequencing data^36^ and a bioinformatic tool previously demonstrated to be reliable for large-scale profiling of STRs^23,35^.

In this work, we leveraged the recently generated expression^5^ and STR^36^ data from 207 wild *C. elegans* strains to conduct a genome-wide scan of how STRs affect gene expression variation. We identified 3,118 and 1,857 expression STRs (eSTRs) that were associated with expression of nearby and remote genes, respectively. We found that eSTRs might help explain missing heritability in SNV-based eQTL studies for both local and distant eQTL. We also explored specific mechanisms of eSTRs and illustrated how local eSTRs might have influenced alternative splicing sites to cause differential transcript usage. We showed that expression of several genes in the same pathway might be altered because of a distant eSTR in a gene upstream. We also found evidence that expression variation in an antioxidant gene, *ctl-1*, might underlie STR variation across wild *C. elegans* strains. We further determined the positive relationship between endogenous oxidative stress and STR insertions/deletions using three mutation accumulation line panels. Our results demonstrate the systemic influences of eSTRs on gene expression and the potential effects of expression variation in antioxidant genes on STR mutations in *C. elegans*. We reveal the interplay between STRs and gene expression variation and provide publicly available frameworks to associate STRs with variation in gene expression and other complex traits in future studies.

## Results

### Variation in STRs regulates expression in nearby genes

We obtained expression data of 25,849 transcripts^5^ of 16,094 genes and 9,691 pSTRs^36^ across 207 wild *C. elegans* strains. We investigated the effects of pSTRs on transcript expression of nearby genes using a likelihood-ratio test (LRT) to evaluate the association between STR variation and transcript expression variation for all pSTRs within 2 Mb surrounding each transcript and with at least two common alleles (allele frequency > 0.5). We applied the LRT using both pSTR genotypes and lengths treating them as factorial variables (See Methods). In total, using STR genotypes, 1,555,828 tests were performed to test the effect of 3,335 pSTRs on the expression variation of 25,849 transcripts, each of which was tested for a median of 59 STRs (ranging from one to 141) (Fig. 1a). Using STR lengths, 1,227,485 tests were performed for the effect of 2,607 pSTRs on the expression variation of 25,847 transcripts, each of which was tested for a median of 47 STRs (ranging from one to 119) (Fig. 1a). For each test, we also performed another test using permuted STR genotypes or lengths. We identified local eSTRs with LRT values that passed the Bonferroni threshold (3.2E-8 and 4.1E-8 for STR genotypes and lengths, respectively) and found 3,082 eSTRs for 2,888 transcripts by STR genotypes and 2,391 eSTRs for 2,791 transcripts by STR lengths, including 2,355 eSTRs for 2,695 transcripts by both STR genotypes or lengths (Fig. 1a, Supplementary Data 1). Each transcript had a median of nine eSTRs (ranging from one to 77) and six eSTRs (ranging from one to 65) by STR genotypes and lengths, respectively. None of the tests using permuted STRs passed the Bonferroni thresholds (Fig. 1a, Supplementary Data 1). As expected, we observed that STRs in close proximity to or within a transcript were more likely to pass the significance threshold than STRs far away from the transcript (Fig. 1a), indicating a close relationship between STRs and gene expression.

**Fig. 1:**
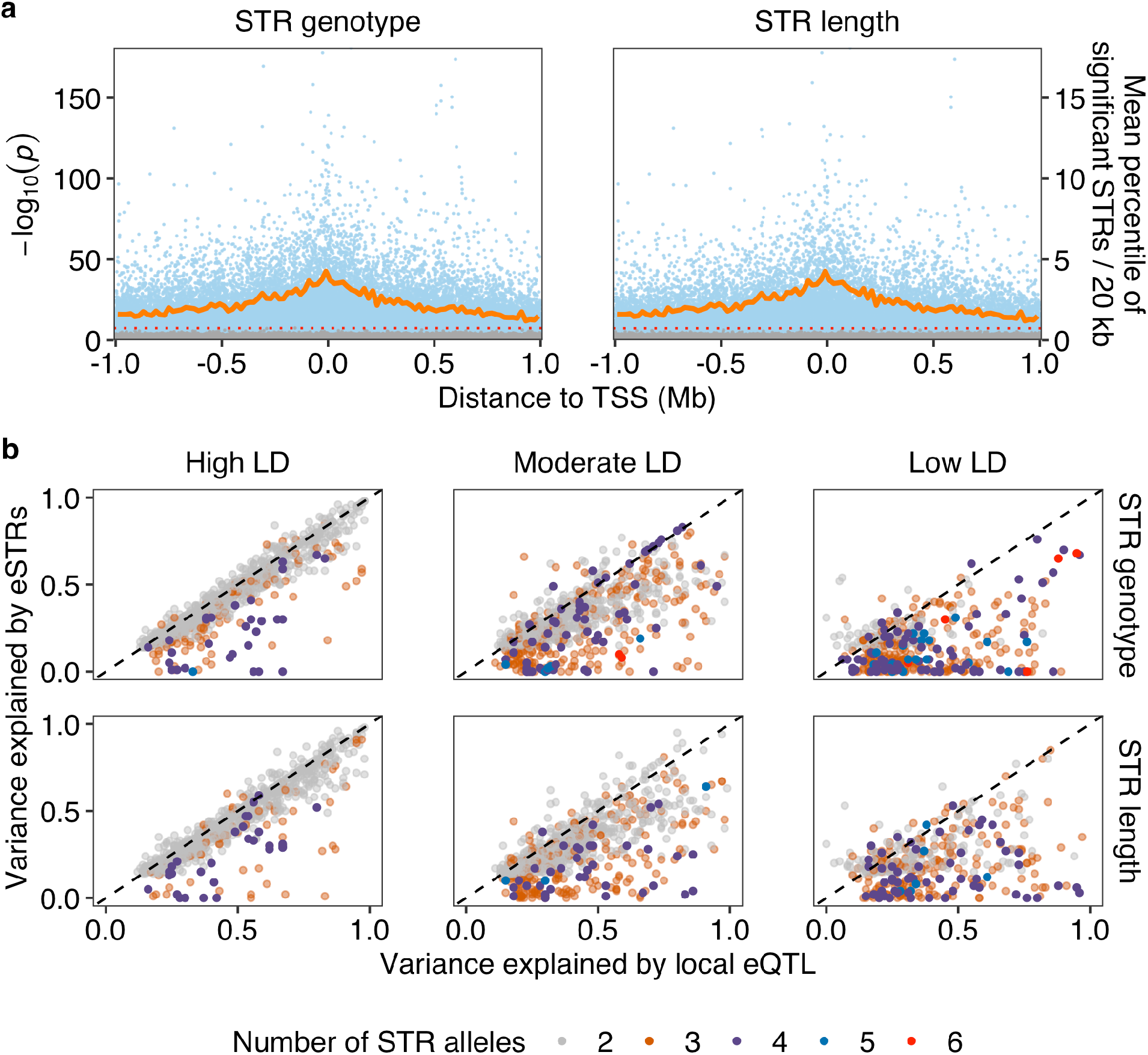
Expression STRs identified using Likelihood-Ratio Tests. **a** Identification of expression STRs (eSTRs) using Likelihood-Ratio Tests (LRT) on full (including STR variation as a variable) and reduced (excluding STR variation as a variable) models. The effects of STR variation in genotype (left panel) or length (right panel) were analyzed separately as factorial variables. Each dot represents a test between STR and transcript expression variation and is plotted with the distance of the STR to the transcription start site (TSS) of the transcript (x-axis) against its -log10(*p*) value (y-axis on the left). Blue and gray dots represent tests using real and permuted data of STR variation, respectively. The red dotted horizontal lines represent Bonferroni thresholds. The dark orange lines represent the mean percentage of significant tests (real data) above the Bonferroni thresholds in each 20 kb bin (y-axis on the right). **b** The variance explained (VE) by local eQTL that were identified using GWA mapping experiments^5^ was plotted against the VE for the most significant eSTRs. Dots are colored by the number of STR alleles used in eSTR VE calculation. LD (*r^2^*) between eQTL and eSTRs were used to separate panels on the x-axis, with high LD (*r^2^* ≥ 0.7), moderate LD (0.3 ≤ *r^2^* <0.7), and low LD (*r^2^* < 0.3). The dashed lines on the diagonal are shown as visual guides to represent VE_eQTL_ = VE_eSTRs_.

In our recent eQTL study^5^, we classified eQTL into local eQTL (located close to the genes that they influence) and distant eQTL (located farther away from the genes that they influence). Among the 3,185 transcripts with local eQTL, 2,477 were also found with eSTRs (Enrichment tested by one-sided Fisher’s Exact Test, with *p* = 2.2E-16). To compare the effects of eQTL and eSTRs in gene regulation, we compared the expression variance explained by eQTL and the most significant eSTR for each transcript and the LD between them (Fig. 1b). Most eQTL-eSTR pairs (48%) with high LD (*r^2^* ≥ 0.7) explained similar levels of expression variance (Fig. 1b), suggesting that these eSTRs might be detected because of the high LD to eQTL or *vice versa*. Among eQTL-eSTR pairs with moderate LD (0.3 ≤ *r^2^* <0.7, 35%) or low LD (*r^2^* < 0.3, 17%), most eQTL explained more variance than eSTRs (Fig. 1b), suggesting these eSTRs, in particular multiallelic eSTRs, might be independent from the eQTL. Although eQTL identified using single-marker based GWA mappings explained a fraction of the variance in gene expression, eSTRs might help explain some missing heritability^37^.

### Insertion in a local eSTR affects transcript isoform usage

We next focused on eSTRs that were in genomic features of their target transcripts and were outside of hyper-divergent regions^38^. We predicted the functional consequences^39^ of these eSTRs and found a total of 13 eSTRs in 16 transcripts of 12 genes that showed high-impact mutations, including missense mutations, in-frame insertions and deletions, start lost, stop gain, and mutations in splicing regions or acceptors. Another 17 eSTRs in 21 transcripts of 17 genes were predicted to affect 5’UTRs and 3’UTRs. We identified two enriched motif sequences, ATTTTT and ATGTT, in these eSTRs by STR genotypes (one-sided Fisher exact test, Bonferroni-corrected *p* = 0.04 and 6.8E-5, respectively) or STR lengths (one-sided Fisher exact test, Bonferroni-corrected *p* = 0.03 and 4.6E-5, respectively). Instead of finding multiple eSTRs, the two motif sequences only came from two eSTRs, STR_13795 of (ATTTTT)_5_ and STR_24584 of (ATGTT)_6.2_, each of which was associated with multiple transcripts of the same genes. In particular, STR_24584 was predicted to have high-impact mutations in the splicing regions of four transcripts of the gene, *R07B7.2*, and was associated with their expression variation (Fig. 2). Compared to strains with the reference allele, strains with a 3-bp insertion showed significantly higher expression in the isoforms *R07B7.2[ab]* but significantly lower expression in the isoforms *R07B7.2[cd]* (Fig. 2a). More specifically, the insertion was located at the 3’ splice site in the intron between exon 7 and exon 8 of *R07B7.2[ab]* and at the junction of the intron and exon 8 for *R07B7.2[cd]* (Fig. 2b). We speculated that at least two mechanisms might underlie the expression differences among the four transcripts caused by STR_24584 variation. First, the insertion [ATT] changed the 3’ splice site of *R07B7.2[ab]* from 5’-GTAACAG-3’ to 5’-TTAACAG-3’ (Fig. 2b), which became closer to the conserved consensus sequence 5’-UUUUCAG-3’ of the 3’ splice site in *C. elegans*^40^. Therefore, the insertion might promote splicing efficiency for *R07B7.2[ab]* in pre-mRNAs of *R07B7.2* and thus increase the expression of the two transcripts, which consequently would decrease the expression of *R07B7.2[cd]*. Second, the insertion could cause a frameshift and insertion in the coding regions of *R07B7.2[cd]*, which caused I474NL (ATA to AATTTA) and V471DL (GTA to GATTTA) in *R07B7.2[c]* and *R07B7.2[d]* (Fig. 2b), respectively. These mutations might increase mRNA degradation. Taken together, our results demonstrated the effects of STR variation on gene expression and provided examples for potential underlying mechanisms.

**Fig. 2:**
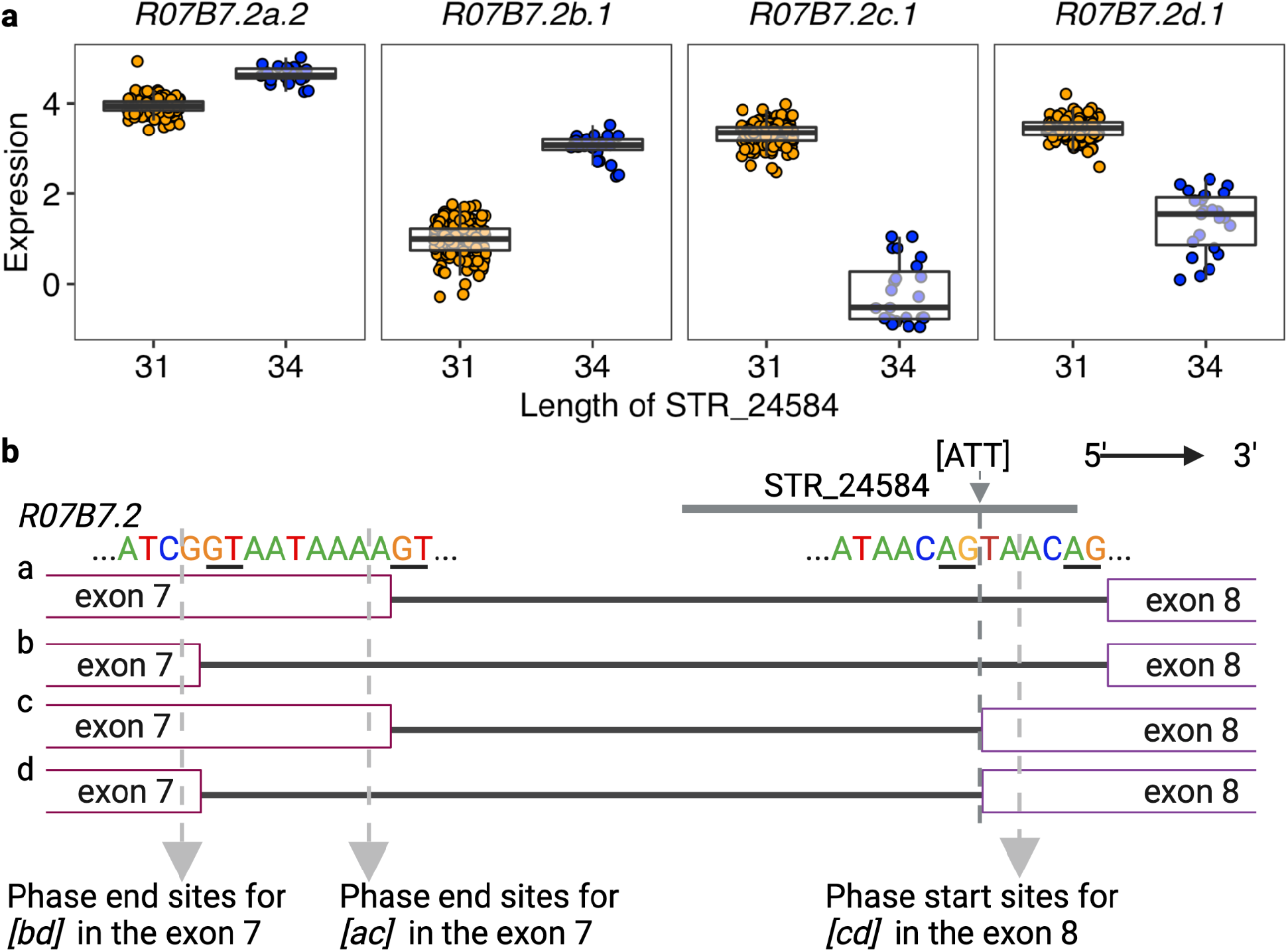
Expression STRs disrupting splicing. **a** Tukey box plots showing expression variation of four transcripts of the gene *R07B7.2* between strains with different lengths of the STR_24584. Each point corresponds to a strain and is colored orange or blue for strains with the N2 reference allele or the alternative allele, respectively. Box edges denote the 25th and 75th quantiles of the data; and whiskers represent 1.5× the interquartile range. **b** Graphic illustration of sequences in the splice site of four transcripts of the gene *R07B7.2* and the position of STR_24584. The dashed arrow in dark gray indicates the position of a 3-bp insertion in the STR_24584 and the splicing region of *R07b7.2[ab]*. The dashed arrow in light gray indicates the phase start and end sites for different exons. Created using BioRender.com.

### STR variation underlies distant eQTL hotspots

In addition to local eQTL, we also identified 3,360 distant eQTL for 2,553 transcripts from 2,382 genes^5^. Genetic variants underlying distant eQTL might affect genes encoding diffusible factors like TFs to regulate genes across the genome. After the identification of local eSTRs, we identified distant eSTRs that affect remote genes. Instead of testing all pSTRs across the genome for each transcript, we selected pSTRs that are within 2 Mb surrounding the QTL regions of interest for all distant eQTL of each transcript. We used LRT tests (as above, also see Methods) to associate pSTR length variation with expression variation. In total, 353,694 tests were performed for the effects of 2,743 pSTRs on the expression variation of 2,553 transcripts, each of which was tested for a median of 104 STRs (ranging from one to 1,005). We used the Bonferroni threshold (1.4E-7) to identify 1,857 distant eSTRs for 950 transcripts, with a median of three distant eSTRs (ranging from one to 127) (Supplementary Data 2). We also compared the expression variation explained by each distant eQTL and the most significant distant eSTR, and the LD between them. Different from local eQTL-eSTR pairs (Fig. 1b), most distant eQTL-eSTR pairs showed moderate (38%) or low (34%) LD, suggesting a more independent role of distant eSTRs in gene regulation (Fig. 3a). We have previously identified 46 distant eQTL hotspots that were enriched with distant eQTL^5^ (Fig. 3b). Genetic variants in these hotspots were associated with expression variation in up to 184 transcripts^5^. Here, we found 229 common distant eSTRs that were associated with at least five distant eQTL in each hotspot (Fig. 3b). Common eSTRs might even underlie about half of all the distant eQTL in several hotspots (Fig. 3b). Altogether, these results suggested the complementary regulatory effects of distant eSTRs to distant eQTL and hotspots.

**Fig. 3:**
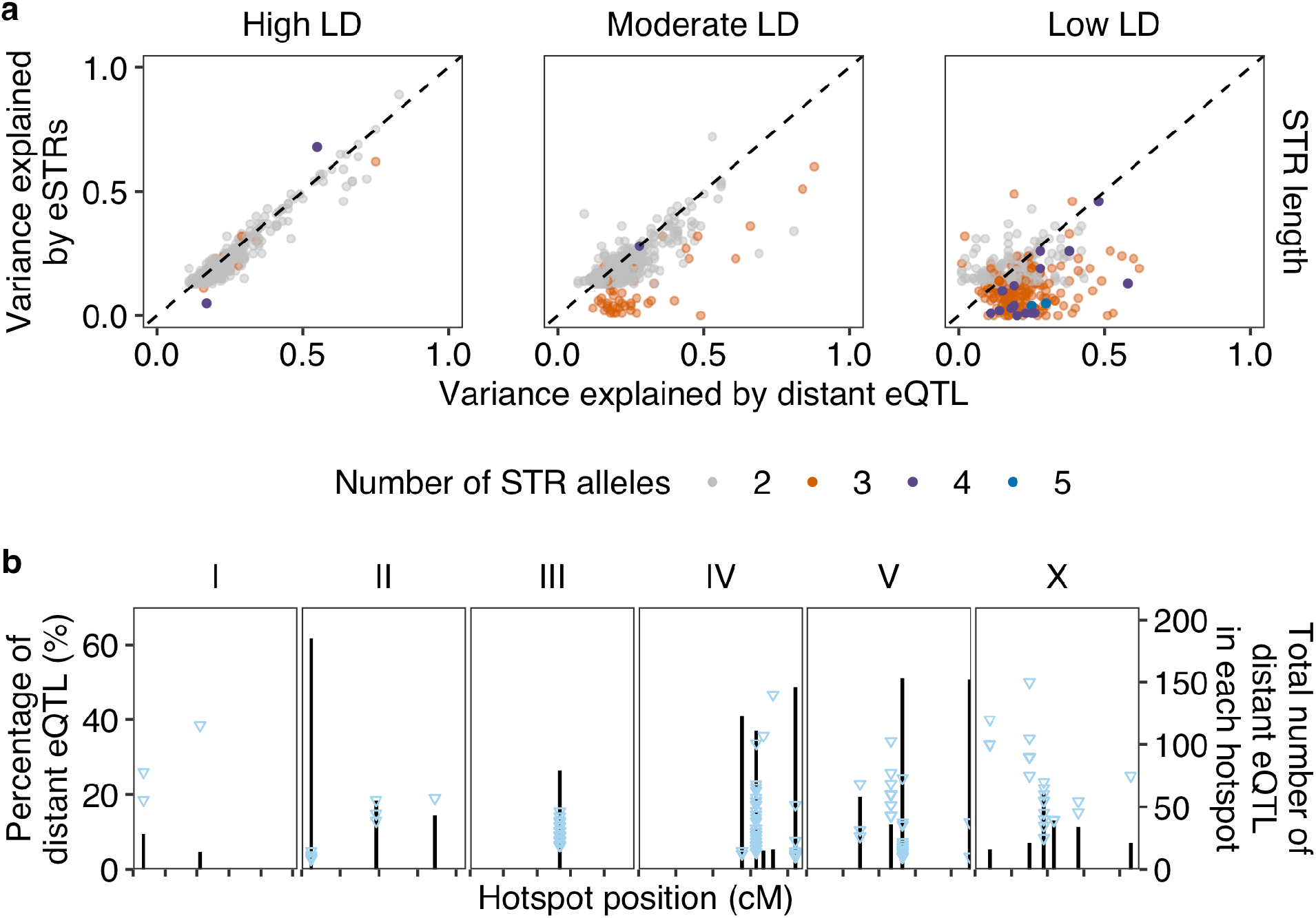
Expression STRs underlying distant eQTL hotspots. **a** The variance explained (VE) by distant eQTL that were identified by GWA mapping experiments^5^ was plotted against the VE by the most significant eSTRs. Dots are colored by the number of STR alleles used in eSTR VE calculation. LD (*r^2^*) between eQTL and eSTRs were used to separate panels on the x-axis, with high LD (*r^2^* ≥ 0.7), moderate LD (0.3 ≤ *r^2^*<0.7), and low LD (*r^2^* < 0.3). The dashed lines on the diagonal are shown as visual guides to represent VE_eQTL_ = VE_eSTRs_. **b** The percentage of distant eQTL (y-axis on the left) that were associated with eSTRs in each distant eQTL hotspot^5^ across the genome (x-axis) is shown. Each blue triangle represents a common eSTR. Black bar indicates the total number of distant eQTL (y-axis on the right) in each hotspot. Tick marks on the x-axis denote every 10 cM.

We next investigated whether any of the common distant eSTRs were in genes encoding TFs or chromatin cofactors. We found nine TF genes and one chromatin cofactors genes that harbor common distant eSTRs (Supplementary Data 3). For example, STR_12763 was a common eSTR for seven distant eQTL in the hotspot ranging from 26 to 27.5 cM on chromosome III (Supplementary Data 3). STR_12763 is in the 3’UTR of the TF gene, *atf-7*^41^, and overlaps with the binding sites of multiple miRNAs (Supplementary Fig. 1). Variation in STR_12763 could affect the targeting of *atf-7* mRNAs by miRNAs to alter expression of the six transcripts (genes). However, none of the ten common distant eSTRs were also identified as local eSTRs for the genes in which they are located. So, we investigated whether any other common eSTRs, although not in known regulatory genes, were also identified as local eSTRs.

We found ten common distant eSTRs that were also local eSTRs for seven genes (Supplementary Data 3). We previously mentioned STR_13795 (ATTTTT)_5_ as one of the two local eSTRs with enriched motif sequences. The variation of STR_13795 was associated with two transcripts of the gene, *cls-2*. Strains with STR contraction by about three repeats (17 bp) in STR_13795 showed significantly higher expression in both transcripts of *cls-2* than strains with the reference STR allele (Supplementary Fig. 2a). Because STR_13795 was in the 3’UTR of *cls-2*, the 17-bp deletion associated with expression of *cls-2* might affect targeting by miRNAs^42,43^. STR_13795 was also identified as a distant eSTR for another ten transcripts, including the gene *polq-1* (Supplementary Fig. 2b). STR_13083 was identified as a local eSTR for *polq-1* and distant eSTRs for another nine transcripts, of which six had STR_13795 as an eSTR (Supplementary Fig. 2b, Supplementary Fig. 3). Most strains with length 30 and 13 in the STR_13795 also have length 16 and 15, respectively in the STR_13083 (Supplementary Table 1). Because STR_13795 was also associated with *polq-1*, STR_13795 was more likely to be the causal candidate than STR_13083 to alter the expression of the six overlapped target transcripts. The significant association between STR_13083 length variation and the expression variation of the six overlapped transcripts were identified because of the linkage between STR_13083 and STR_13795. The three transcripts that only had STR_13083 as their distant eSTRs could also be associated with the length variation of STR_13795, which was not tested for the three transcripts because it was too distant from the genes. Altogether, STR_13795 might affect the expression of all the 13 remote transcripts and genes by altering the expression of *cls-2* (Supplementary Fig. 2, Supplementary Fig. 3b). We performed gene set enrichment analysis for the 13 genes on WormBase^44^ and found significant enrichment in genes related to spindle and germline defectiveness (Supplementary Table 2). The conserved protein, CLASP/CLS-2, is required for mitotic central spindle stability, oocyte meiotic spindle assembly, chromosome segregation, and polar body extrusion in *C. elegans*^45–49^. To summarize, variation in STR_13795 might alter the expression of *cls-2*, which could further affect other related genes in the spindle assembly pathways.

### Oxidative stress potentially drives STR mutations

To explore the genome-wide influences of STRs on gene expression variation, we also wondered what factors might affect STR mutations and cause STR variation across *C. elegans*. DNA strand slippage during replication, DNA repair, and recombination processes can lead to STR mutations^24^. We reasoned that any genetic or environmental factors that are able to increase errors during these processes or decrease genome stability could increase STR mutation rates^50,51^. We hypothesized that, if variation in genetic factors that affect genomic stability exists, the amount of total STR variation could be used as a quantitative trait for a GWA mapping study. We recently also developed a pipeline of mediation analysis to link gene expression variation to quantitative traits^5^. Thus, we sought to examine potential genetic and mediating factors underlying STR mutation variation.

We first defined an STR variation trait by counting reference and alternative STR alleles for each of the 207 strains in the 9,691 pSTRs (See Methods) (Supplementary Fig. 4a). Deletions are the predominant mutations in STR mutations across wild *C. elegans* strains (Supplementary Fig. 4a). We performed GWA mappings using two methods, LOCO and INBRED^52^, for this trait (see Methods). The INBRED method corrects more heavily for genetic stratification and many times decreases mapping power more than the LOCO method^52–54^. We detected six QTL with large QTL regions of interest on five of the six chromosomes using LOCO but no QTL using INBRED (Supplementary Fig. 4b, Supplementary Table 3). We next used mediation analysis to link expression differences with total STR mutation variation. Mediation analysis was performed for any transcripts with eQTL that overlap with the QTL regions of interest of the six QTL for STR variation. We identified 31 significant mediator transcripts of 26 genes (Fig. 4a). The top mediator gene, *ctl-1*, had two transcripts identified as significant mediators by multiple tests using different pairs of eQTL and QTL (Fig. 4a). We found moderate negative correlations between the expression of the two *ctl-1* transcripts (*Y54G11A.6.1* and *Y54G11A.6.2*) and STR mutation variation (Fig. 4b), suggesting that the expression level of *ctl-1* might impact STR mutation variation. We regressed the STR variation trait by the expression of the transcript *Y54G11A.6.1* and performed GWA mappings. All the QTL mapped using the raw trait and LOCO disappeared in the mappings using the regressed trait (Supplementary Fig. 4c, Supplementary Table 3), supporting that the expression variation of *ctl-1* might affect STR mutation variation. We also identified a new QTL at the position 14,625,147 on chromosome II in both LOCO and INBRED methods (Supplementary Fig. 4c, Supplementary Table 3), suggesting that loci other than *ctl-1* might affect STR mutation variation as well.

**Fig. 4:**
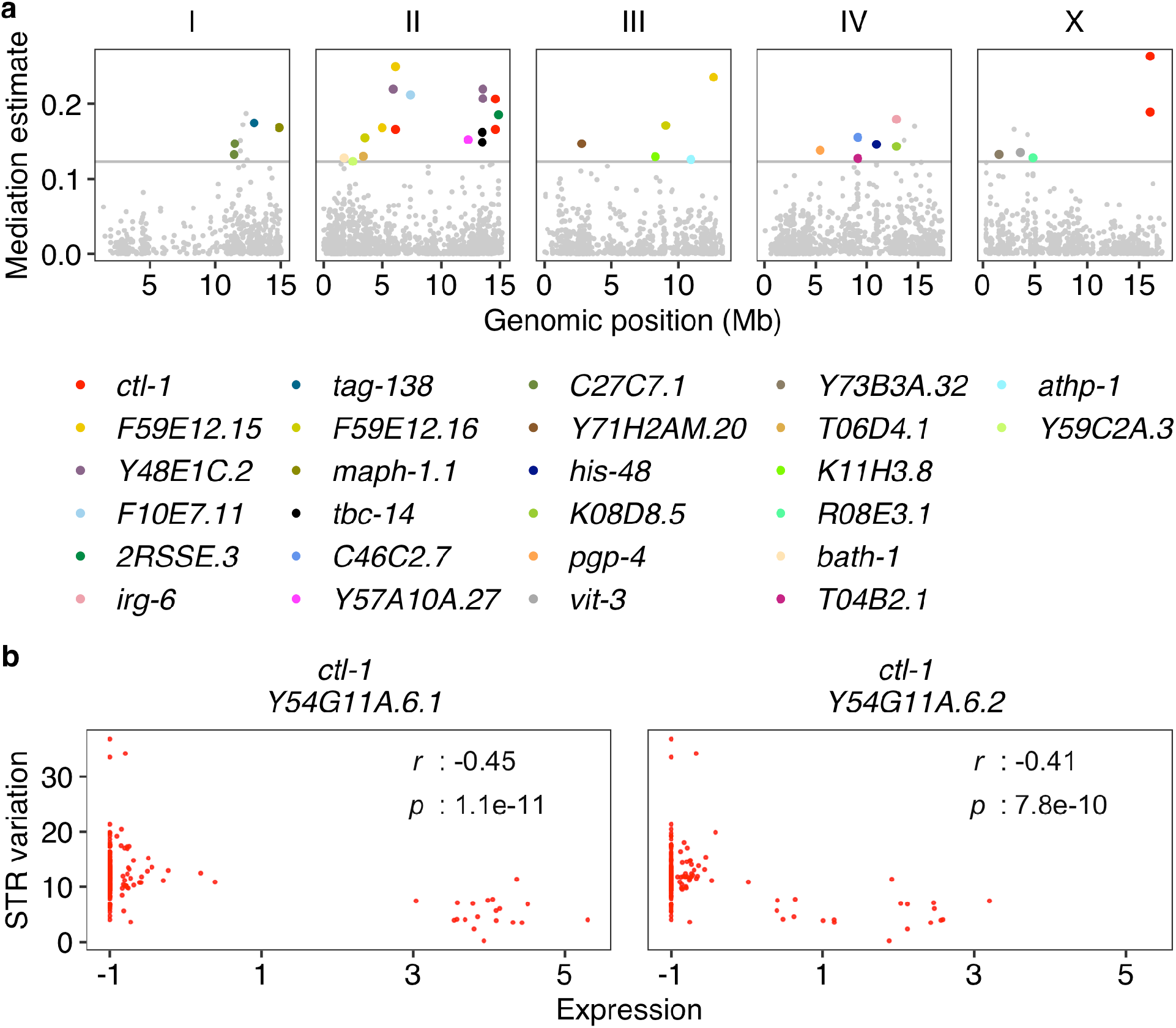
Mediation effects of *ctl-1* expression on STR variation. **a** Mediation estimates (y-axis) of transcript expression on STR variation are plotted against the genomic position (x-axis) of the eQTL. The horizontal gray line represents the 99^th^ percentile of the distribution of mediation estimates. Mediator transcripts with adjusted *p* < 0.05 and interpretable mediation estimate greater than the 99^th^ percentile estimates threshold are colored by their genes. Other tested mediator transcripts are colored gray. **b** The correlation of expression (x-axis) of four mediator transcripts to STR variation (y-axis) is shown. Each dot represents a strain and is colored by mediator genes as in **a**. The coefficient *r* and the *p*-value for each correlation using the two-sided Pearson’s correlation tests are indicated in the top right.

The gene, *ctl-1*, encodes a cytosolic catalase in the detoxification pathway of reactive oxygen species (ROS)^55^. Elevated expression of *ctl-1* and other antioxidant related genes, which likely enhanced resistance to oxidative stresses, was associated with lifespan elongation in *C. elegans*^56,57^. Oxidative damage can alter DNA secondary structure, affect genome stability and replication, and cause mutations^58^. Therefore, it is possible that the group of strains showing high levels of *ctl-1* expression managed to reduce STR mutations caused by oxidative damage over time and have lower levels of total STR mutations across the species (Fig. 4b). We have previously detected five (one local and four distant) and six (one local and five distant) eQTL for expression variation of the two transcripts of *ctl-1, Y54G11A.6.1* and *Y54G11A.6.2*, respectively^5^. Among the 5,291 transcripts with detected eQTL, 4,430 transcripts had a single eQTL detected and only 30 transcripts were found with equal or more than five eQTL^5^. These results suggest that the expression of *ctl-1* was highly controlled and might be critical for adaptation to oxidative stresses.

We further examined potential relationships between oxidative stresses and STR mutations using three mutation accumulation (MA) line panels^59–62^ that have undergone passage for many generations with minimal selection: 1) 67 MA lines that were derived from N2 and propagated for ~250 generations; 2) 23 MA lines that were derived from a mutant strain, *mev-1* (with a missense mutation introgressed into N2, resulting in elevated oxidative stress), and propagated for ~125 generations; and 3) 67 MA lines that were derived from PB306 (a wild strain) and propagated for ~250 generations. We obtained raw sequencing data for these 157 MA lines and their three ancestors and called STR variation using the same method that we used for wild *C. elegans* strains^36^ (See Methods). We calculated mutation rates for three different mutations (deletions, insertions, and substitutions) between the ancestor and each derived MA line and compared mutation rates across the three MA lines (Fig. 5). We found that *mev-1* MA lines showed significantly higher mutation rates in deletions and insertions but significantly lower substitution rates than the other two MA lines (Fig. 5, Supplementary Table 4). The gene *mev-1* encodes a mitochondrial complex II SDHC^63^. The *mev-1* mutant was found to be highly sensitive to oxidative stress and showed reduced lifespan^63^. The high deletion and insertion rates in *mev-1* lines might be driven by their increased endogenous oxidative damage than the other two MA lines. Although the mutation rate of substitution was low in *mev-1* lines, deletions and insertions likely contributed most of the variation in STRs (Supplementary Fig. 4A).

**Fig. 5:**
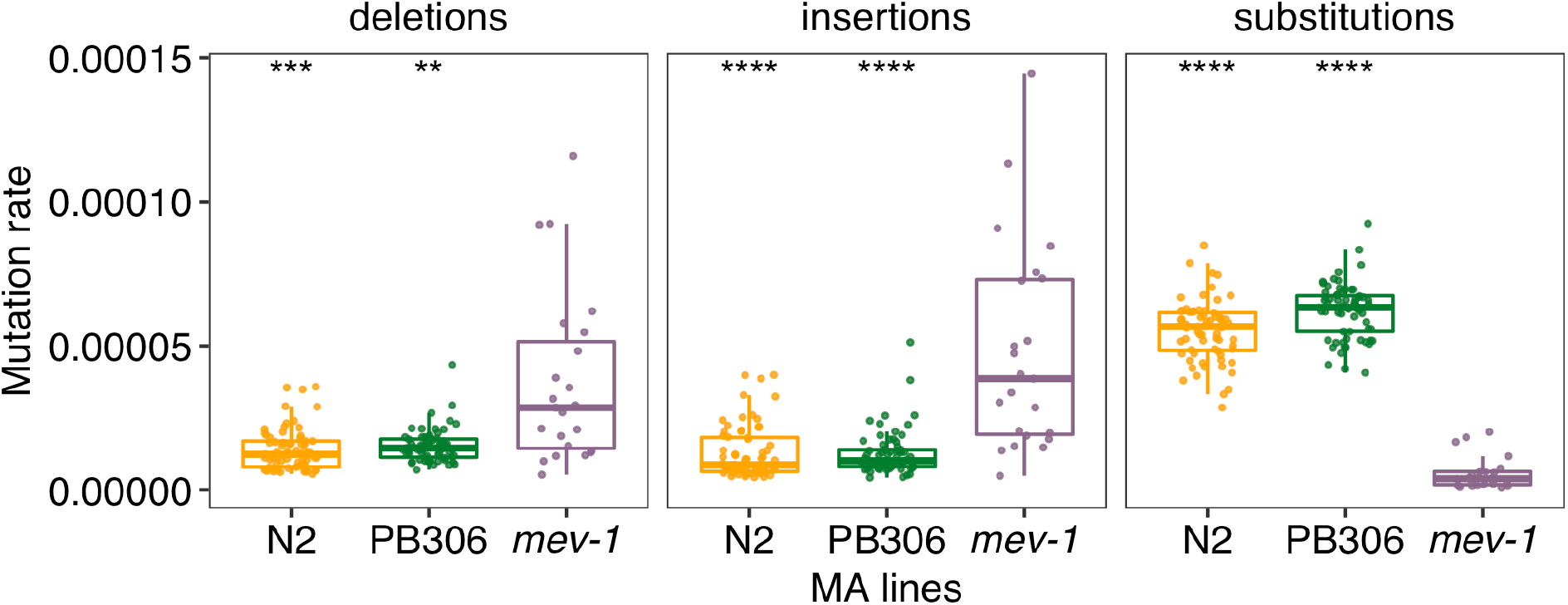
STR mutation rates in the MA lines. Comparison of STR mutation rates in deletions, insertions, and substitutions between the *mev-1* line (purple) and N2 (orange), and PB306 (green) lines, respectively. Box edges denote the 25th and 75th quantiles of the data; and whiskers represent 1.5× the interquartile range. Statistical significance of difference comparisons (Supplementary Table 4) was calculated using the two-sided Wilcoxon test and *p*-values were adjusted for multiple comparisons (Bonferroni method). Significance of each comparison is shown above each comparison pair (**: adjusted *p* ≤ 0.01; ***: adjusted *p* ≤ 0.001; ****: adjusted *p* ≤ 0.0001).

Altogether, these results suggest that oxidative stresses affect variation in STRs. Although a laboratory mutation in *mev-1* might have increased oxidative stresses and led to more deletions and insertions in STRs, natural genetic variation that promoted the expression of *ctl-1* might reduce oxidative stress, which might stabilize STRs to prevent mutations (Fig. 4b).

## Discussion

Natural allelic variation in different classes of genomic loci contributes to gene expression variation^3–5,17–19^. We previously identified thousands of eQTL correlated with SNVs across wild *C. elegans* strains^5^. Here, we performed genome-wide analysis on how one of the most polymorphic and abundant repetitive elements, STRs^36^, might affect expression variation in *C. elegans*. We identified nearly 5,000 associations between STR variation and expression variation of nearby and remote genes (Fig. 1, Fig. 3). It is important to note that the number of eSTRs that we detected only represents a conservative estimate because of the strict significance threshold that we applied.

We previously performed genome-wide association analysis on phenotypic variation in 11 organismal complex traits using pSTR length variation^36^ and SNVs^64–73^ respectively. Most of the significant STRs were located within or close to the QTL regions of interest identified using SNVs and GWA mappings, indicating close relationships between significant STRs and QTL. In the detection of eSTRs, we modeled pSTRs^36^ within 2 Mb surrounding each of the 25,849 transcripts with reliable expression data^5^ (Fig. 1). Close to 84% of transcripts found with local eSTRs were previously detected with local eQTL^5^, indicating close relationships between eSTRs and eQTL. Therefore, we further modeled pSTRs within 2 Mb surrounding the QTL regions of interest for transcripts with detected distant eQTL. Our results revealed important roles of distant eSTRs underlying distant eQTL and hotspots (Fig. 3). Among transcripts with both eSTRs and eQTL, 48% of local and 28% of distant eSTR-eQTL pairs showed strong LD with each other and explained similar amounts of expression variance (Fig. 1b, Fig. 3a). Future work using simulations and experiments is necessary to partition the contributions of eSTRs and eQTL to gene regulatory differences. Additionally, we also found 17% of local and 34% of distant eSTR-eQTL pairs showed low LD with each other (Fig. 1b, Fig. 3a). Among these low LD eSTR-eQTL pairs, 69% of local and 60% of distant eSTRs had three to six alleles used in LRT tests (Fig. 1b, Fig. 3a), indicating independent roles of eSTRs, especially multiallelic STRs, in explaining expression variance. Note that the LD between eQTL and multiallelic STRs might be overestimated because we transformed multiallelic STR genotypes to biallelic to calculate LD (See Methods). Therefore, potentially more multiallelic eSTRs than we reported could have affected expression independently from eQTL, which could help explain the missing heritability in complex gene expression traits. One future direction that we did not explore is how epistasis, the interactions between STRs and SNVs or other mutations, affects gene expression^74,75^.

STRs have been proposed to regulate gene expression using various molecular mechanisms^17,25,76–79^. We found local eSTR variants that caused a variety of mutations in the target transcripts. We dissected how a 3-bp insertion in an eSTR of the gene *R07B7.2* altered 3’ splice site to change alternative splicing efficiency and cause differential transcript usage (Fig. 2). The function of the gene *R07B7.2* is not well understood but the expression of *R07B7.2* was found enriched in neurons, such as AVG and RIM^44^. Future efforts could investigate the neural consequences of different transcript usage in the gene *R07B7.2*. Furthermore, we found that distant eSTRs might affect gene expression by disrupting miRNA binding in the 3’UTRs of genes encoding TFs, such as ATF-7 (Supplementary Fig. 1). Although the variation of STR_12763 and expression variation of *atf-7* was not significant in the local eSTR identification, it is possible that the effects of STR_12763 variation on the expression of *atf-7* were too small to be detected using data from 207 strains. But the small changes in the abundance of the ATF-7 protein might cause strong expression differences in the ATF-7 targets, which were detectable within the power of our study. In addition to TFs, we also identified that the eSTR STR_13795 might affect four genes (*cls-2, ddx-23, pck-2*, and *F54E7.9*) in the spindle assembly pathways through both local and distant regulation. It is possible that *cls-2* is at the upstream of the pathway and its expression could affect the other three downstream genes. Several mutants of *cls-2* have been generated^80^. Future work could use these mutants to first examine whether the expression of *cls-2* affects the other three genes and then validate the role of STR_13795 mutations in expression regulation.

Not only did we observe eSTR that altered gene expression, we also found that gene expression variation might affect STR mutations. We performed GWA mappings and mediation analysis on an STR variation trait and identified a candidate gene, *ctl-1*, that functions in the detoxification pathway of reactive oxygen species (ROS) (Fig. 4, Supplementary Fig. 4b). We observed low levels of genome-wide STR mutations in strains with high expression of *ctl-1* (Fig. 4b), which might have increased the antioxidant capacity in the animal to stabilize the genome and reduce mutations. The effects of ROS on STR mutations were also revealed by *mev-1* MA lines, which experienced elevated oxidative stresses and showed higher STR deletion and insertion rates than wild type MA lines (Fig. 5).

Not every strain with low levels of STR mutations had high levels of *ctl-1* expression (Fig. 4b), suggesting STR mutations are polygenic. For example, other genes that are responsible for stress response in *C. elegans* might also affect STR mutations. Fungal infections were found to induce STR expansion in wheat^50^. Various natural pathogens of *C. elegans* have been discovered^81–84^, future work could compare STR mutations among *C. elegans* strains isolated from locations with or without known pathogens. Additionally, genes that are related to DNA replication, repair, or the mitotic process, such as the second top mediator, *F59E12.15* (Fig. 4a), could also cause genome-wide effects on STR mutations.

Altogether, our study provides the first large-scale analysis of associations between STRs and gene expression variation in wild *C. elegans* strains. We highlighted the role of eSTRs in explaining expression variation and missing heritability. We also proposed that oxidative stress might have driven STR mutations globally. STRs have been proposed to facilitate adaptation and accelerate evolution^12,16–19,25,31,85^. Future work could use our data and analysis framework to study how STR variation affects complex traits and facilitates adaptation of *C. elegans* in the wild.

## Methods

### *C. elegans* expression and STR data

We obtained summarized expression data of 25,849 transcripts of 16,094 genes and genotypes of 9,691 polymorphic STRs (pSTRs) in 207 *C. elegans* strains from the original studies^5,36^. We also obtained 6,545 eQTL positions, their QTL regions of interest, and eQTL classification from the *C. elegans* eQTL study^5^.

### Expression STRs (eSTRs) identification

#### STR genotype transformation

Genotypes of each pSTRs for each strain were transformed as previously described^36^. Briefly, we used single digits (*e.g*., “0”, “1”, “2”) to represent STR genotypes in strains with homozygous alleles (*e.g*., “0|0”, “1|1”, “2|2”); we chose the smaller digits (*e.g*., “0”, “1”, “2”) to represent STR genotypes in strains with heterozygous alleles (*e.g*., “0|1”, “1|2”, “3|2”).

#### Selection of STRs for eSTRs identification tests

To identify local eSTRs, we selected pSTRs within 2 Mb surrounding each of the 25,849 transcripts with reliable expression measurements^5^. To identify distant eSTRs, we selected pSTRs within 2 Mb surrounding the QTL regions of interest for each of the 2,553 transcripts with detected distant eQTL^5^. Among selected pSTRs for each transcript, we further selected STRs with at least two common variants (frequency > 0.05) among strains with both STR genotype and expression data, and only retained strains with common STR variants.

#### Likelihood-ratio test (LRT) to identify eSTRs

We treated STR genotypes as factorial variables and performed LRT on the full model *lm(expression ~ STR*) and the reduced model *lm(expression ~ 1*) using the *lrtest(*) function in the R package *lmtest* (v0.9-39) (https://cran.r-project.org/web/packages/lmtest/index.html). The Bonferroni threshold was used to identify significant eSTRs. For each test using real data, we also performed another LRT using permuted data by shuffling STR genotypes across strains.

#### eSTR identification using STR length variation

Because different alleles of the same STR might have the same length and STR length variation might have stronger effect on gene expression than substitution, we performed LRT using the mean allele length of the two copies of each STR for each strain as factorial variables. We performed STR selection, permutation, LRT, and the Bonferroni threshold as above to identify eSTRs using STR length variation.

### LD and variance explained by eQTL and eSTRs

We calculated linkage disequilibrium (LD) between top eSTRs and eQTL for transcripts with both regulatory sites detected. We used eQTL genotypes and STR genotypes to calculate LD for eSTRs detected by both STR genotype variation and STR length variation. Only strains used in eSTR identification were used for LD calculation. We acquired genotypes of wild strains at each eQTL from the hard-filtered isotype variant call format (VCF) file (CeNDR 20210121 release)^86^. For processed STR genotypes, we further transformed all multiallelic variants into biallelic variants by converting all non-reference genotypes (1,2,3, etc.) to 1 and kept reference genotypes as 0. Then, we calculated LD correlation coefficient *r^2^* for each STR-SNV and SNV-SNV pairs using the function *LD()* in the R package *genetics* (v1.3.8.1.2) (https://cran.r-project.org/package=genetics). We also used the generic function cor() in R and Pearson correlation coefficient to calculate the expression variance explained by each QTL and each top eSTR.

### Genetic basis of STR variation

#### STR variation trait

We performed GWA mapping to identify the genetic basis of STR variation in *C. elegans*. For each of the 207 strains, we counted the total number of STRs with no missing genotypes among the 9,691 polymorphic STRs and the total number (*N_total_*) of alternative alleles (*N_alt_*) for both copies at each site. The STR variation trait, which is used as the phenotypic input of GWA mappings, was calculated as *log_10_*(*N_alt_*/*2N_total_*).

#### Genome-wide association (GWA) mappings

We performed GWA mappings using the pipeline *Nemascan* (https://github.com/AndersenLab/NemaScan) as previously described^52^. Briefly, we extracted SNVs of the 207 strains from the hard-filtered isotype VCF (CeNDR 20210121 release)^86^ and filtered out variants that had any missing genotype calls and variants that were below the 5% minor allele frequency using *BCFtools* (v.1.9)^39^. We further pruned variants with a LD threshold of *r^2^* ≥ 0.8 using *-indep-pairwise 50 10 0.8* in *PLINK* (v1.9)^87,88^ to generate the genotype matrix containing 20,402 markers. We then used two approaches in the software *GCTA* (v1.93.2)^53,54^ to perform GWA mappings: 1) the leave-one-chromosome-out (LOCO) approach, which uses the *-mlma-loco* function to both construct a kinship matrix using variants in all chromosomes except the chromosome in testing and perform the GWA mapping; and 2) the INBRED approach, which uses the *-maker-grm-inbred* function to construct a kinship matrix that is designated for inbred organisms and the *-fastGWA-lmm-exact* function for the GWA mapping^52–54^. An eigen-decomposition significance threshold (EIGEN) and a more stringent Bonferroni-corrected significance threshold (BF) were estimated in *Nemascan* for QTL identification. For EIGEN, we first estimated the number of independent tests (*N_test_*) within the genotype matrix using the R package *RSpectra* (v0.16.0) (https://github.com/yixuan/RSpectra) and *correlateR* (0.1) (https://github.com/AEBilgrau/correlateR). EIGEN was calculated as *-log_10_(0.05/N_test_*). BF was calculated using all tested markers. Here, QTL were defined by at least one marker that was above BF. QTL regions of interest were determined by all markers that were above BF and within 1 kb of one another, and 150 more markers on each flank.

#### Mediation analysis

We performed mediation analysis that is implemented in *Nemascan* to identify the mediation effect of gene expression on STR variation as previously described^5^. Briefly, for each QTL of STR variation, we used the genotype (*Exposure*) at the QTL, transcript expression traits (*Mediator*) that have eQTL^5^ overlapped with the QTL, and STR variation (*Outcome*) as input to perform mediation analysis using the *medTest()* function in the R package *MultiMed* (v2.6.0) (https://bioconductor.org/packages/release/bioc/html/MultiMed.html) and the *mediate()* function in the R package *mediation* (v4.5.0)^89^. Significant mediators were identified as those with adjusted *p* < 0.05 and interpretable mediation estimate greater than the 99^th^ percentile of all estimates.

#### GWA mapping for the regressed STR variation trait

We regressed the STR variation trait by the expression of the transcript *Y54G11A.6.1* of the gene *ctl-1* and performed GWA mappings as described above.

### STR variants in mutation accumulation (MA) lines

We obtained whole-genome sequence data in the FASTQ format of 160 MA lines, including N2 MA lines: the N2 ancestor and 67 MA lines; *mev-1* MA lines: the *mev-1* ancestor and 23 MA lines; and PB306 MA lines: the PB306 ancestor and 67 MA lines (NCBI Sequence Read Archive projects PRJNA395568, PRJNA429972, and PRJNA665851)^61,62^. We used the pipelines *trim-fq-nf* (https://github.com/AndersenLab/trim-fq-nf) and *alignment-nf* (https://github.com/AndersenLab/alignment-nf) to trim raw FASTQ files and generate BAM files for each line, respectively^86^. We called STR variants for the 160 lines using the pipeline *wi-STRs* (https://github.com/AndersenLab/wi-STRs)^36^.

### Mutation rate of polymorphic STRs in MA lines

We calculated the STR mutation rate in MA lines as previously described^36^ but using variant calls before filtering by 10% missing data. Briefly, between each MA line and its ancestor, we selected STR sites with reliable (“PASS”) calls in both lines. Then, for each STR, we compared the two alleles in the MA line to the two alleles in the ancestor, respectively, to identify insertion, deletion, substitution, or no mutation. The mutation rate (per-allele, per-STR, per-generation) *μ* for each type of mutation was calculated as *m/2nt* where *m* is the number of the mutation, *n* is the total number of reliable STRs, and *t* is the number of generations^61,62^.

### Statistical analysis

Statistical significance of difference comparisons were calculated using the Wilcoxon test and *p*-values were adjusted for multiple comparisons (Bonferroni method) using the *compare_means()* function in the R package *ggpubr* (v0.2.4) (https://github.com/kassambara/ggpubr/). Enrichment analyses were performed using the one-sided Fisher’s exact test and were corrected for multiple comparisons (Bonferroni method).

## Acknowledgments

We would like to thank Timothy A. Crombie and Ryan McKeown for helpful comments on the manuscript. G.Z. is supported by the NSF-Simons Center for Quantitative Biology at Northwestern University (awards Simons Foundation/SFARI 597491-RWC and the National Science Foundation 1764421). E.C.A. is supported by a National Science Foundation CAREER Award (IOS-1751035) and a grant from the National Institutes of Health R01 DK115690. The *C. elegans* Natural Diversity Resource is supported by a National Science Foundation Living Collections Award to E.C.A. (1930382). We would also like to thank WormBase because without it these analyses would not have been possible.

## Author contributions

E.C.A. and G.Z. designed the study. G.Z. analyzed the data. G.Z. and E.C.A. wrote the manuscript.

## Data availability

The datasets for generating all figures can be found at https://github.com/AndersenLab/Ce-eSTRs. Expression and eQTL data of wild *C. elegans* strains were obtained from https://github.com/AndersenLab/WI-Ce-eQTL^5^. *C. elegans* STR variation data were obtained from https://github.com/AndersenLab/WI-Ce-STRs^36^. The hard-filtered isotype VCF (20210121 release) was obtained from CeNDR (https://www.elegansvariation.org/data/release/20210121). The raw sequencing data of MA lines were obtained from the NCBI Sequence Read Archive under accession code PRJNA395568 [https://www.ncbi.nlm.nih.gov/bioproject/PRJNA395568], PRJNA429972 [https://www.ncbi.nlm.nih.gov/bioproject/PRJNA429972], and PRJNA665851) [https://www.ncbi.nlm.nih.gov/bioproject/PRJNA665851]^61,62^.

## Code availability

The code for generating all figures can be found at https://github.com/AndersenLab/Ce-eSTRs.

## Supplementary Information

### Description of Additional Supplementary Files

File Name: Supplementary Data 1

Description: List of eSTRs for nearby genes.

File Name: Supplementary Data 2

Description: List of eSTRs for remote genes.

File Name: Supplementary Data 3

Description: List of common distant eSTRs that are in genes encoding TFs or chromatin cofactors, or the distant eSTRs are also local eSTRs for the genes in which they are located.

### Supplementary Tables

**Supplementary Table 1.** Number of strains with REF or ALT allele lengths in STR_13795 and STR_13083. Only 186 strains with expression data and genotypes at both STR sites are included.

| STR |  |  | STR_13795 |  |
| --- | --- | --- | --- | --- |
|  | Allele |  | REF | ALT |
|  |  | Length | 30 | 13 |
| STR_13083 | REF | 16 | 133 strains | 15 strains |
|  | ALT | 15 | 6 strains | 32 strains |

**Supplementary Table 2.**
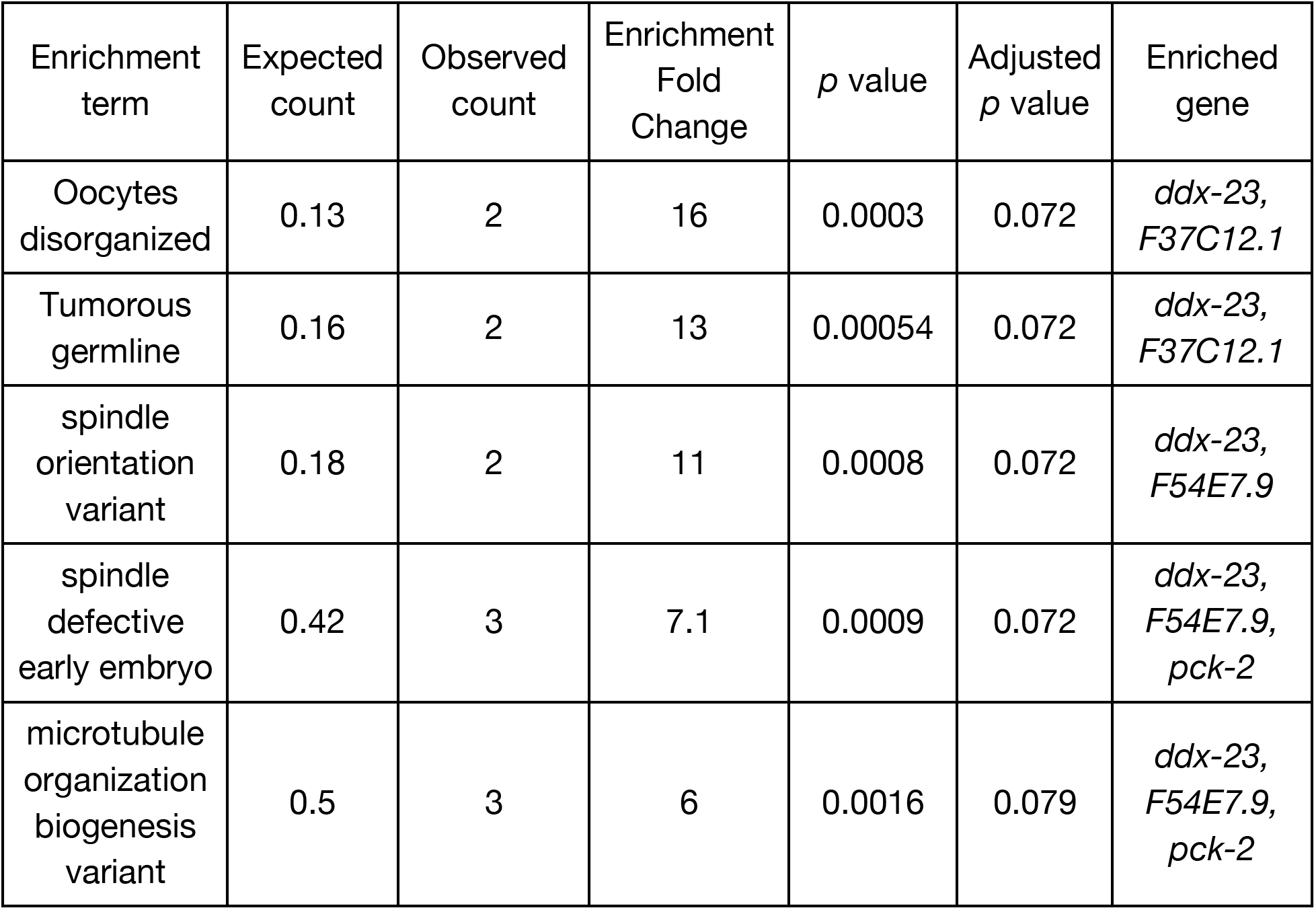
GSEA results of ten genes that were associated with STR_13795 in the gene *cls-2*.

| Enrichment term | Expected count | Observed count | Enrichment Fold Change | *p* value | Adjusted *p* value | Enriched gene |
| --- | --- | --- | --- | --- | --- | --- |
| Oocytes disorganized | 0.13 | 2 | 16 | 0.0003 | 0.072 | *ddx-23*,<br>*F37C12.1* |
| Tumorous germline | 0.16 | 2 | 13 | 0.00054 | 0.072 | *ddx-23*,<br>*F37C12.1* |
| spindle orientation variant | 0.18 | 2 | 11 | 0.0008 | 0.072 | *ddx-23*,<br>*F54E7.9* |
| spindle defective early embryo | 0.42 | 3 | 7.1 | 0.0009 | 0.072 | *ddx-23*,<br>*F54E7.9*,<br>*pck-2* |
| microtubule organization biogenesis variant | 0.5 | 3 | 6 | 0.0016 | 0.079 | *ddx-23*,<br>*F54E7.9*,<br>*pck-2* |

**Supplementary Table 3.** GWA QTL and regions of interest for the raw and regressed STR variation traits.

| Trait | GWA method | QTL chromosome | QTL peak position | Start position of QTL region of interest | End position of QTL region of interest |
| --- | --- | --- | --- | --- | --- |
| STR variation | LOCO | I | 12153609 | 4754801 | 13671576 |
| STR variation | LOCO | II | 2706647 | 1477894 | 15272855 |
| STR variation | LOCO | III | 4892173 | 2807201 | 13718059 |
| STR variation | LOCO | IV | 13760657 | 2278611 | 15378958 |
| STR variation | LOCO | X | 2603542 | 826476 | 8910667 |
| STR variation | LOCO | X | 14551237 | 14217837 | 17696299 |
| Regressed STR variation | LOCO | II | 11566198 | 10505390 | 11842443 |
| Regressed STR variation | LOCO | II | 14625147 | 13968708 | 15272855 |
| Regressed STR variation | INBRED | II | 14625147 | 13968708 | 15272855 |

**Supplementary Table 4.** Comparison of mutation rates among MA lines using two-sided Wilcoxon tests and Bonferroni method for multiple testing correction.

| Mutation | group1 | group2 | *p* value | Adjusted<br>*p* value |
| --- | --- | --- | --- | --- |
| deletions | *mev-1* | N2 | 2.46E-05 | 0.00015 |
| deletions | *mev-1* | PB306 | 1.79E-04 | 0.0011 |
| insertions | *mev-1* | N2 | 1.56E-07 | 9.4E-07 |
| insertions | *mev-1* | PB306 | 1.67E-08 | 1E-07 |
| substitutions | *mev-1* | N2 | 1.06E-12 | 6.3E-12 |
| substitutions | *mev-1* | PB306 | 1.06E-12 | 6.3E-12 |

### Supplementary Figures

**Supplementary Fig. 1.**
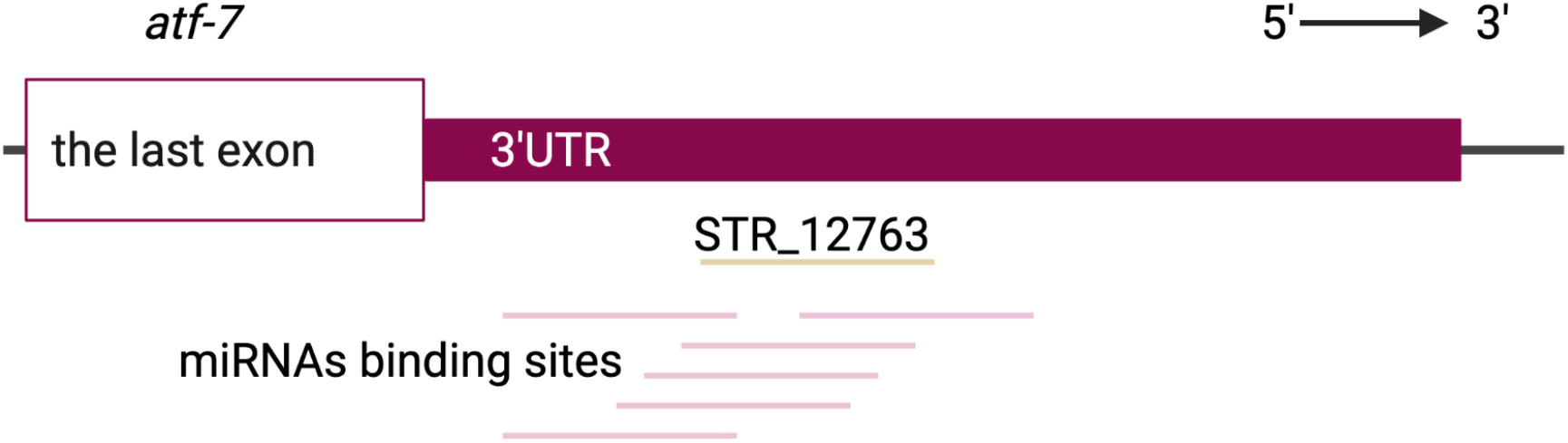
STR_12763 in 3’UTR of the TF gene, *atf-7*, might affect miRNA binding sites. Graphic illustration of the 3’UTR of *atf-7*, the STR_12763 (the light brown line), and predicted binding sites of miRNAs (pink lines) based on WormBase^1^. Created using BioRender.com.

**Supplementary Fig. 2.**
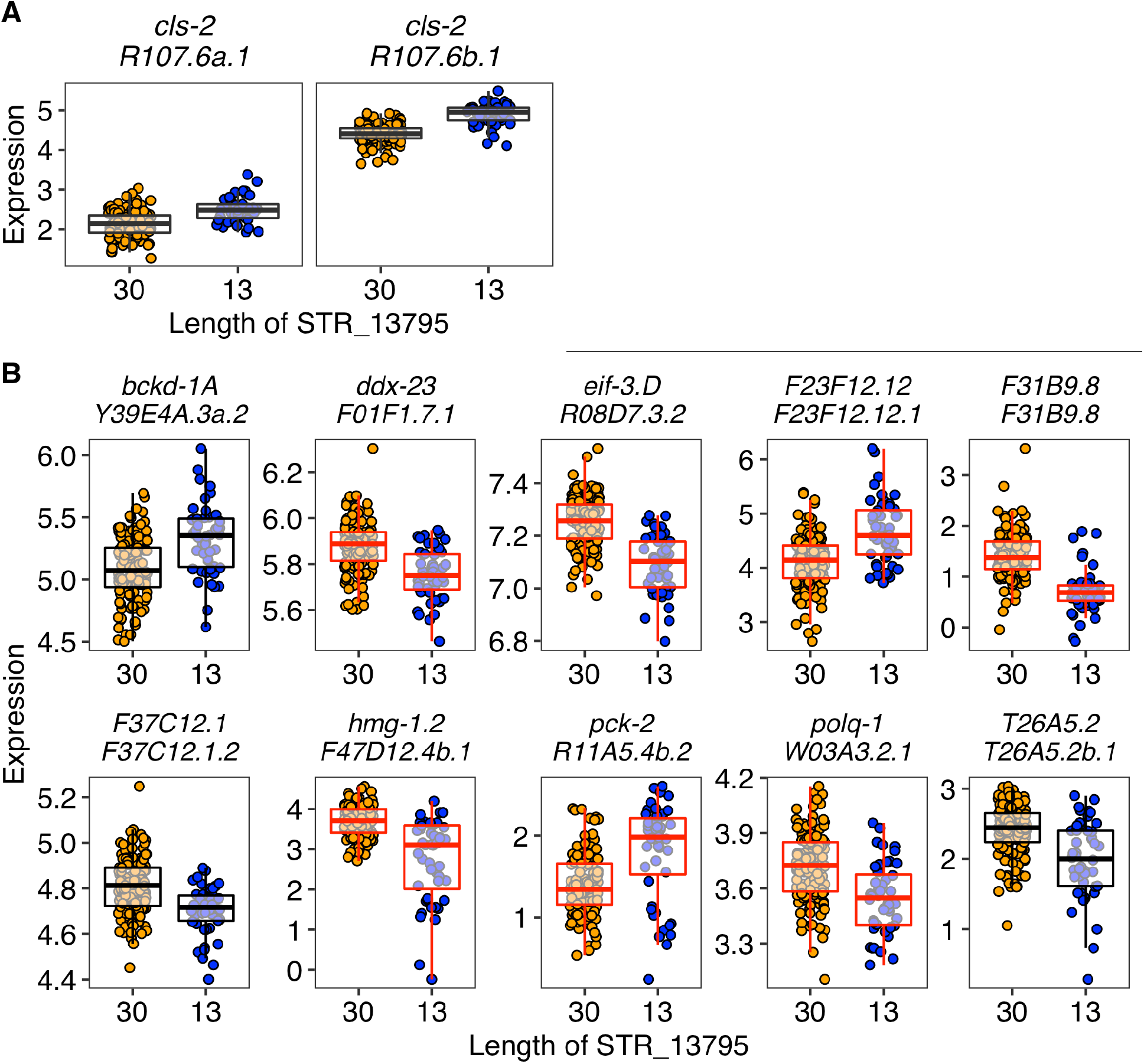
The local and distant eSTR, STR_13795. STR_13795 was identified as local eSTRs for two transcripts of the gene *cls-2* **(A)** and distant eSTRs for ten other transcripts **(B)**. Tukey box plots showing expression variation of the 12 transcripts between strains with different lengths of the STR_13795 are shown and colored red for those transcripts with STR_13083 as an eSTR (Supplementary Fig. 3). Each point corresponds to a strain and is colored orange and blue for strains with the N2 reference allele and the alternative allele, respectively. Box edges denote the 25th and 75th quantiles of the data; and whiskers represent 1.5× the interquartile range.

**Supplementary Fig. 3.**
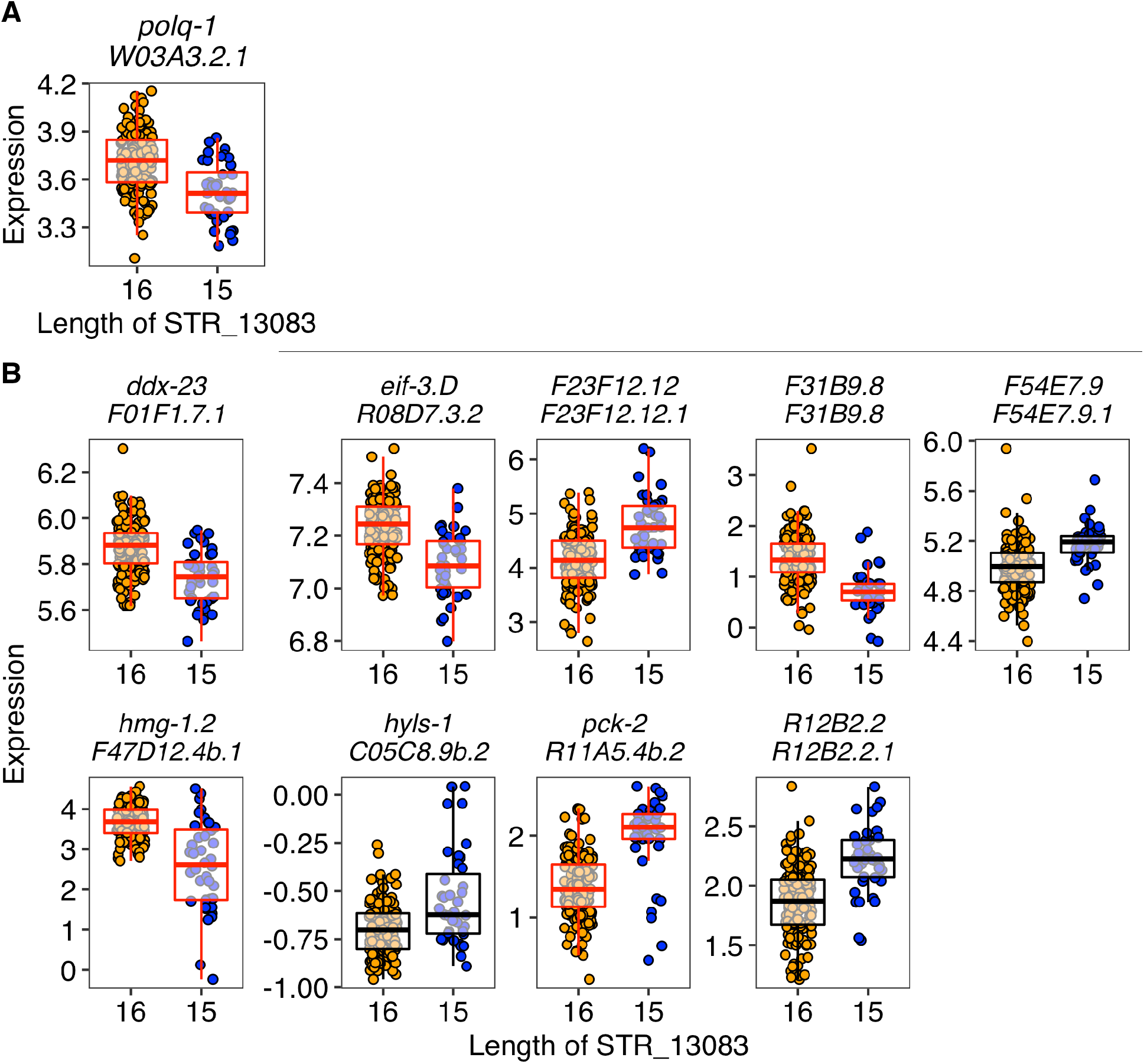
The local and distant eSTR, STR_13083. STR_13083 was identified as local eSTRs for the transcript of the gene *polq-1* **(A)** and distant eSTRs for nine other transcripts **(B)**. Tukey box plots showing expression variation of the ten transcripts between strains with different lengths of the STR_13083 are shown and colored red for those transcripts with STR_13795 as an eSTR (Supplementary Fig. 2). Each point corresponds to a strain and is colored orange and blue for strains with the N2 reference allele and the alternative allele, respectively. Box edges denote the 25th and 75th quantiles of the data; and whiskers represent 1.5× the interquartile range.

**Supplementary Fig. 4.**
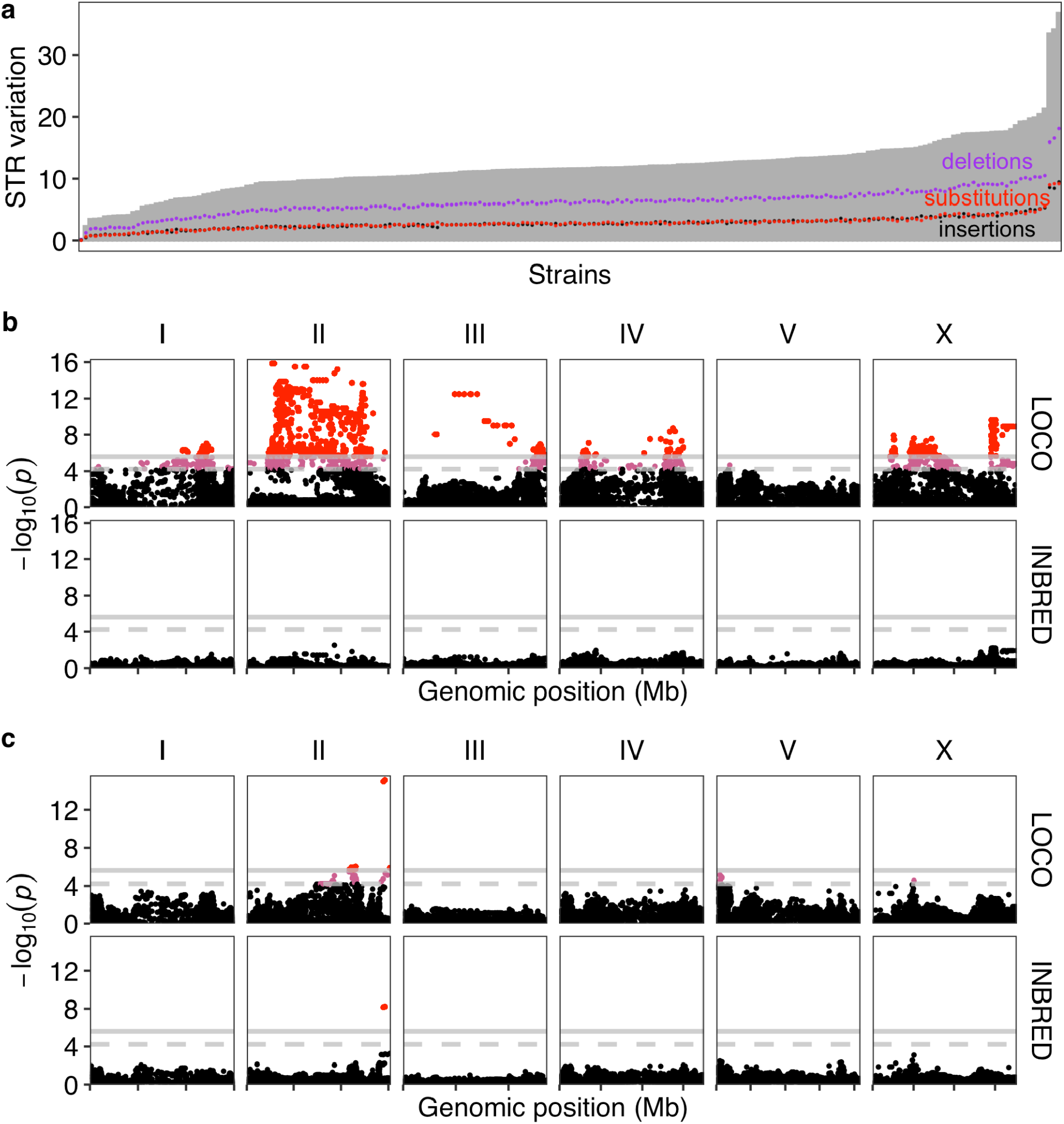
Genetic basis underlying STR variation. **a** The distribution of an STR variation trait across 207 strains is shown. The STR variation traits calculated by deletions, insertions, and substitutions for each strain are shown as dots and colored purple, black, and red, respectively. **b** Manhattan plots indicating the GWA mapping results for STR variation across 207 strains using LOCO and INBRED approaches are shown, respectively. **c** Manhattan plots indicating the GWA mapping results for STR variation regressed by the expression of *Y54G11A.6.1* of the gene *ctl-1* across 206 strains using LOCO and INBRED approaches are shown, respectively. In **b** and **c**, each point represents an SNV that is plotted with its genomic position (x-axis) against its -log10(*p*) value (y-axis) in mapping. SNVs that pass the genome-wide EIGEN threshold (the dashed gray horizontal line) and the genome-wide Bonferroni threshold (the solid gray horizontal line) are colored pink and red, respectively. QTL were identified using the Bonferroni threshold.

